# Sand flies and Toscana virus: intra-vector infection dynamics and impact on *Phlebotomus perniciosus* life-history traits

**DOI:** 10.1101/2024.02.25.582017

**Authors:** Lison Laroche, Anne-Laure Bañuls, Rémi Charrel, Albin Fontaine, Nazli Ayhan, Jorian Prudhomme

**Affiliations:** MIVEGEC, Université de Montpellier – IRD – CNRS, Centre IRD, Montpellier, France; Institute of Environmental Sciences (CML), Leiden University, Leiden, Netherlands; Unité des Virus Emergents (UVE: Aix Marseille Université, IRD 190, Inserm 1207, AP-HM Hôpitaux Universitaires de Marseille) Marseille, France; Unité de virologie, Département Microbiologie et maladies infectieuses, Institut de Recherche Biomédicale des Armées (IRBA), Marseille, France; Centre National de Référence des Arbovirus, Marseille, France; Université de Rennes, Inserm, EHESP, IRSET (Institut de Recherche en Santé Environnement Travail), UMR_S 1085, 35000, Rennes, France

**Keywords:** *Phlebovirus*, Toscana virus, sand fly, infection dynamics, experimental infection, transmission

## Abstract

Toscana virus (TOSV) is a leading cause of summer viral meningitis in central Italy and south of France, and can cause severe neurological cases. Within the Mediterranean basin, it is transmitted by hematophagous sand flies belonging to the *Phlebotomus* genus. Despite the identification of the primary TOSV vectors, the virus’s developmental cycle in vector species remains largely unknown. Limited research has been conducted on transmission dynamics and the vectorial competence and capacity of the principal TOSV vector, *Phlebotomus perniciosus*. In this context, we investigated the intra-vector TOSV infection dynamics in *Ph. perniciosus*, as well as its impact on the vector’s life history traits. Female sand flies were experimentally infected with TOSV though an artificial blood meal. Systemic dissemination of the virus was observed approximately three days post-infection, potentially resulting in a shorter extrinsic incubation period. Moreover, the study revealed a longer hatching time for eggs laid by infected females. This research not only confirmed the vector competence of *Ph. perniciosus* but also provided the first insight into TOSV’s developmental cycle and its impact on the vector. These findings prompt further exploration of TOSV transmission dynamics, raise new hypotheses on the virus transmission and highlight the importance of follow-up studies.

**Author summary:** Toscana virus (TOSV) is a reemerging sandfly-borne virus causing neuroinvasive infections in humans. This virus is endemic in the Mediterranean basin, with a potential risk of introduction in northern Europe and Asia. Despite decades of research, few studies have focused on the development cycle of TOSV in sand flies and the dynamics of transmission. Here, we provide a comprehensive study of the intra-vector dynamics of TOSV infection and its impact on both vector biology and transmission. Through experimental infections of the major vector *Phlebotomus perniciosus*, we not only confirmed vector competence but also provided the first insight into the TOSV developmental cycle in the vector by estimating the extrinsic incubation period at six days. Our study reveals an impact of TOSV infection on vector hatching time leading to a delayed emergence of infected sand flies, with a potential impact on transmission. Our findings encourage further exploration of transmission dynamics, raise new hypotheses on alternative transmission pathways, and emphasize the importance of follow-up studies.

## Introduction

Toscana virus (TOSV) is an enveloped, protein-encapsidated, tri-segmented RNA virus that belongs to the *Phlebovirus toscanaense* species, *Phlebovirus* genus, *Phenuiviridae* family and Bunyavirales order (1). Initially isolated in 1971 from *Phlebotomus perniciosus* and *Phlebotomus perfiliewi* in central Italy, TOSV is endemic in the Mediterranean basin where at least 250 million people are at risk of infection (2,3). Despite the increasing frequency of TOSV infections, the virus remains largely neglected due to inadequate diagnostic tools (4). As typical with arthropod-borne viruses (arboviruses), many TOSV infections are unreported since they are either asymptomatic or cause only mild symptoms (5). Toscana virus has a particular neurotropism and is one of the primary causes of meningitis and encephalitis in endemic regions (6). Human infections occur during warm seasons, with a peak in the hottest months in relation to vector activity (3). Three genetic lineages (A, B and C) of TOSV circulate within the Mediterranean area, but no differences in pathogenicity have been demonstrated to date. Toscana virus is transmitted to humans through a bite of an infected female sand fly (7) and its geographical distribution is related to sand fly presence and continues to expand. Currently, four sand fly species are recognized or suspected as TOSV vectors: *Ph. perniciosus*, *Ph. perfiliewi*, *Phlebotomus sergenti* and *Phlebotomus neglectus* (8). However, other sand fly species may also contribute to the virus maintenance and transmission (8).

Due to its neuroinvasive nature, TOSV has emerged as the most significant sandfly-borne phlebovirus for public health, warranting further investigation into its natural cycle (9). However, to date, limited information is available on natural cycle and transmission of TOSV particularly with regard to the phlebovirus infection dynamics in sand flies (10–12). While transovarial and venereal transmissions have been experimentally observed in its primary vector, *Ph. perniciosus* (6), their significance in the natural TOSV cycle remains uncertain and warrants further investigation. Recently it was hypothesized that transmission from infected to uninfected sand flies during sugar feeding might play a significant role in the virus natural cycle (13). This mode of transmission has already been shown experimentally with Massilia virus, another phlebovirus genetically related to TOSV (14). Moreover, TOSV persistence and infectivity in sand fly sugar meal has recently been demonstrated in laboratory conditions (13). Nevertheless, the relevance of this alternative transmission route to TOSV cycle in natural ecosystems remains to be confirmed.

In recent decades, arboviruses have emerged as significant causes of death and disability worldwide, highlighting the need for a better understanding of their transmission dynamics (15–17). Arbovirus epidemics are influenced by both intrinsic factors (vector competence, virus strain or genetic lineage, virus dose-effect, etc.) and extrinsic factors (temperature, rainfall, human activity, etc.) that can influence vector biology (18–20). These factors, which vary temporally and spatially, can affect the infection and transmission dynamics of arboviruses, including TOSV (21). Moreover, certain studies have already examined arbovirus infection impact on vector life-history traits and shown effects on survival and fecundity, with a potential impact on transmission dynamics (22–24). Therefore, to comprehensively address the public health impact of arboviruses, such as TOSV, further studies are needed to examine their infection dynamics.

In our study, we aim to understand the impact of TOSV infection on biology of its primary vector *Ph. perniciosus* and its potential effect on transmission. We set up experimental infections to achieve two main objectives. The first one is to determine the infection dynamics of TOSV in *Ph. perniciosus* by confirming the vector competence for TOSV lineage B and explore how different TOSV dose influence the kinetics of infection within its vector. The second objective is to examine the impacts of TOSV infection on the vector life history traits, including survival and fecundity, and the effects on transmission dynamics. Ultimately, this work will provide a deeper understanding of TOSV transmission mechanisms and the impact of infection on vector biology.

## Results

### Toscana virus infection dynamics

A total of 1836 female sand flies were used to determine the infection dynamics of TOSV in *Ph. perniciosus*. The number of sand flies used in each group depended on the productivity of the sand fly colony at the beginning of the experiment. Respectively, 73 (20%), 761 (68%) and 335 (77%) of females fed on TOSV infected blood with dose 1 (10^2^ TCID_50_/ml), dose 2 (10^4^ TCID_50_/ml) and dose 3 (10^6^ TCID_50_/ml). However, for dose 2 infection monitoring was extended to 17 and 21 days post infection (dpi) as more blood-fed females were available. After blood feeding, sand flies were dissected every two days, however due to the daily natural mortality (10-20%), it was not possible to dissect all blood-fed females. The dissected number of females are summarized in **Table 1**.

**Table 1.** Number of *Phlebotomus perniciosus* dissected per day for Toscana virus infection depending on viral loads.

|  | Day post infection |  |  |  |  |  |  |  |  |
| --- | --- | --- | --- | --- | --- | --- | --- | --- | --- |
|  | 3 | 5 | 7 | 9 | 11 | 13 | 15 | 17 | 21 |
| Dose 1 (10<sup>2</sup> TCID<sub>50</sub>/ml) | 10 | 5 | 5 | 6 | 5 | 5 | 2 | ∅ | ∅ |
| Dose 2 (10<sup>4</sup> TCID<sub>50</sub>/ml) | 20 | 47 | 39 | 46 | 52 | 55 | 45 | 26 | 30 |
| Dose 3 (10<sup>6</sup> TCID<sub>50</sub>/ml) | 20 | 20 | 20 | 20 | 10 | 10 | 4 | ∅ | ∅ |

After testing sand flies: head, body, wings and legs, at dose 1, TOSV was detected by RT-qPCR in the bodies of two out of 10 females at three dpi and one out of five females at seven dpi. No virus was found in heads or wings and legs after infection with dose 1. For the dose 2 infection, we observed viral load higher than 10^4^ RNA copies/ml at three dpi in female bodies. This viral load increased then to 10^6^ RNA copies/ml at 11 dpi, indicating viral replication (**Fig 1, A**). The viral load in the heads, wings and legs was lower compare to bodies, but started to increase from four to five dpi. We observed a stabilization of the viral load in the wings and legs from nine dpi, remaining between 10^2^ and 10^3^ RNA copies/ml until 15 dpi. In sand flies infected with dose 2, viral load continued to increase to reach 2.10^6^ (body), 8.10^4^ (head) and 2.10^4^ (wings and legs) RNA copies/ml up to 21 days (data not shown). For the infection with dose 3, we detected TOSV RNA in all sand fly parts (head, body, wings and legs) until 15 dpi (**Fig 1, B**). The viral load increased by almost one log from three to 13 dpi in bodies. The viral load started to increase in the head, wings and legs from four to five dpi, but with a lower RNA copy number. The analysis of significance of the TOSV dose on post-infection day showed that the viral load has a significant effect on the infection dynamics in sand flies (*P* = 0.0002) (data from dose 1 infection was not included in the analysis due to low sample size). The incidence of infection exhibited a dose-dependent augmentation in response to TOSV exposure, with a significant effect of post-infection time (*P* = 0.0314).

**Figure 1.**
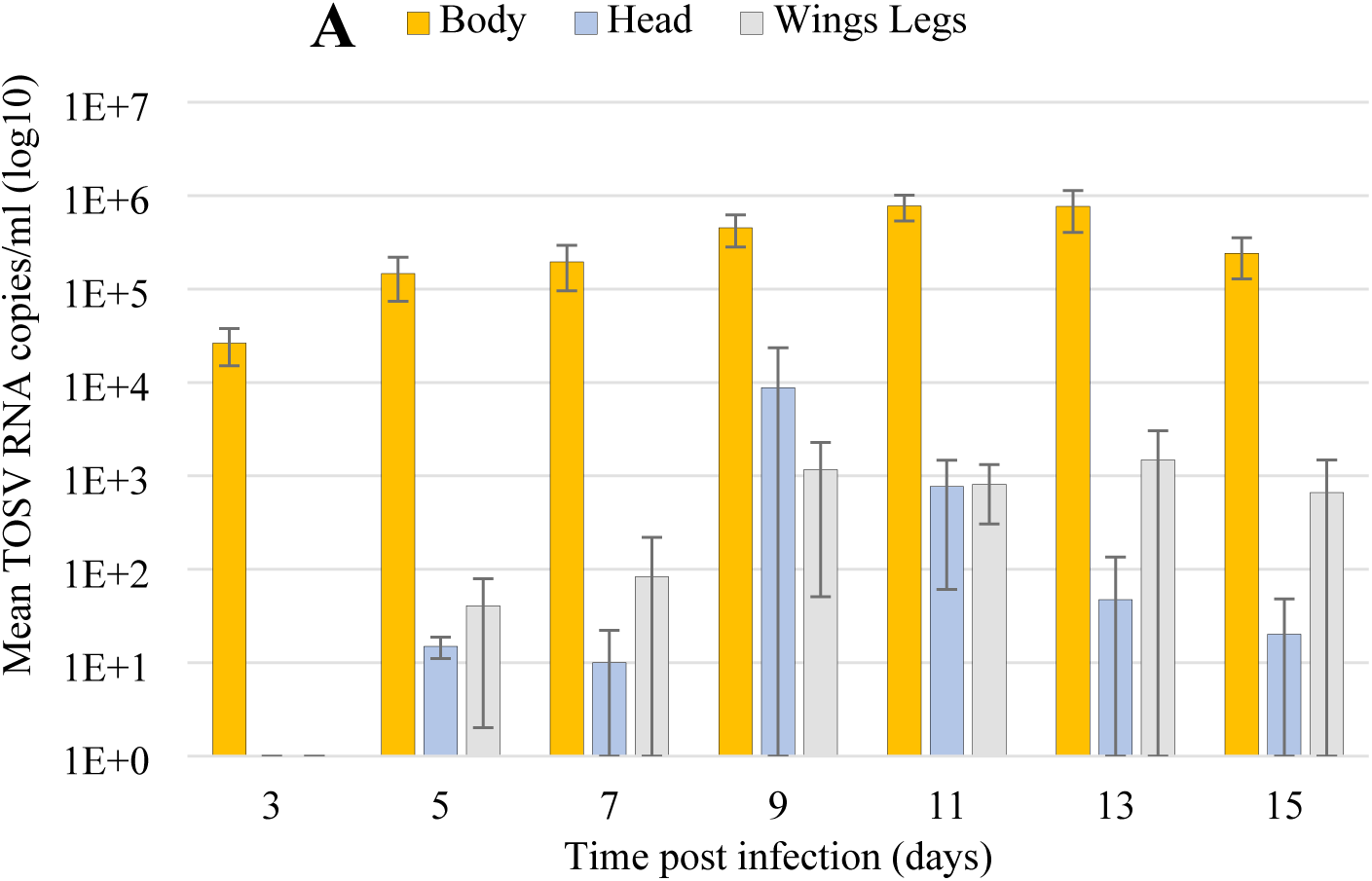
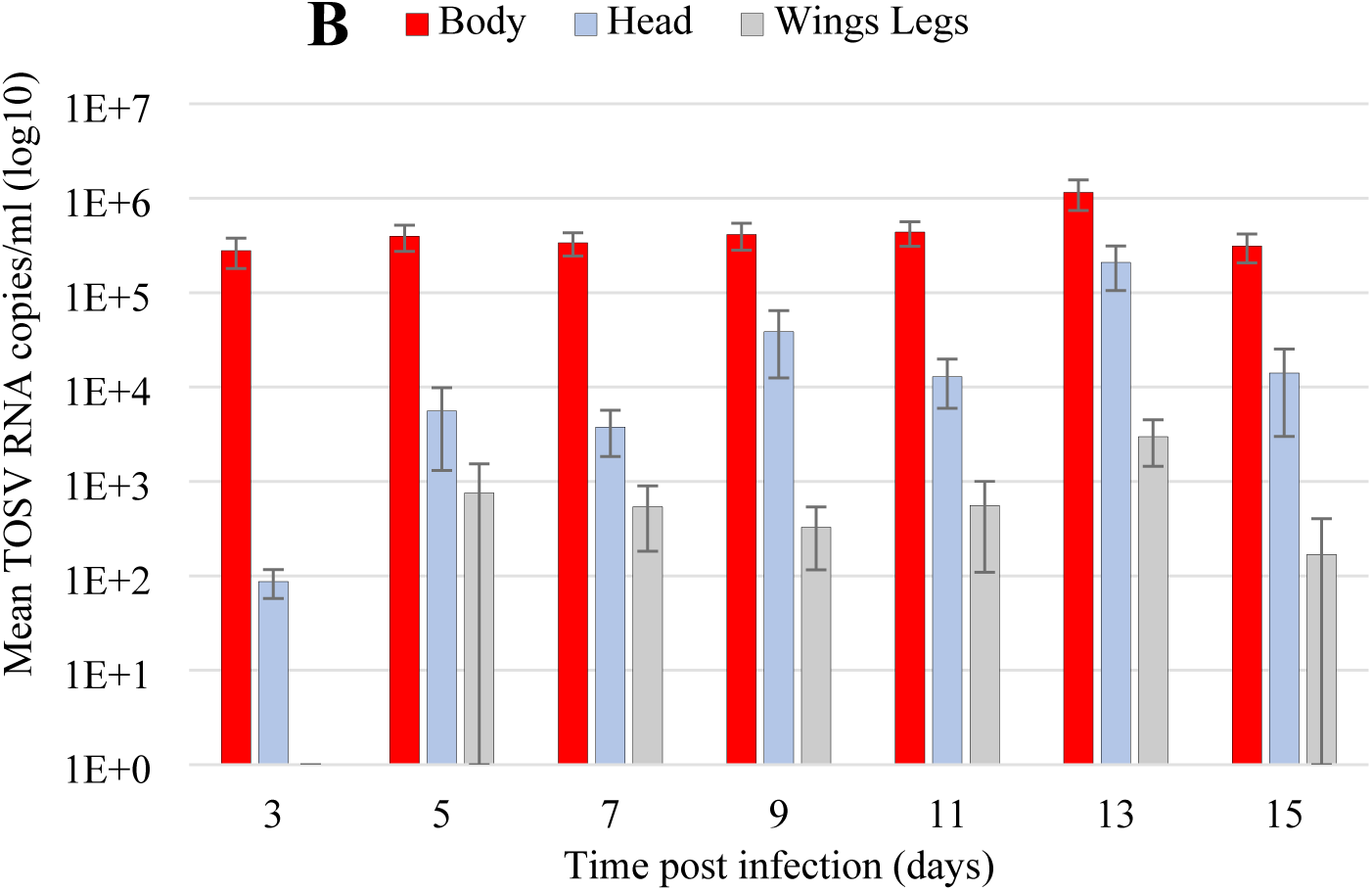
Toscana virus RNA copies/ml (log10) in body, head, wings and legs of infected females with dose 2 (A) and dose 3 (B) of Toscana virus over time.

Systemic infection rates were measured by comparing the number of sand flies with infected heads to infected bodies. The within-host dynamics of systemic TOSV infection were represented for the two viral doses by fitting a non-linear regression model (**Fig 2**). The rates of systemic infection did not saturate at 100% for either TOSV dose. The estimated time of 50% systemic infection was approximately five (SE: 0.1) days for sand flies infected with dose 2. For sand flies infected with dose 3, the estimated time of 50% systemic infection was three (SE: 0.3) days, but due to the low effective numbers, this estimate can be considered between zero and five.

**Figure 2.**
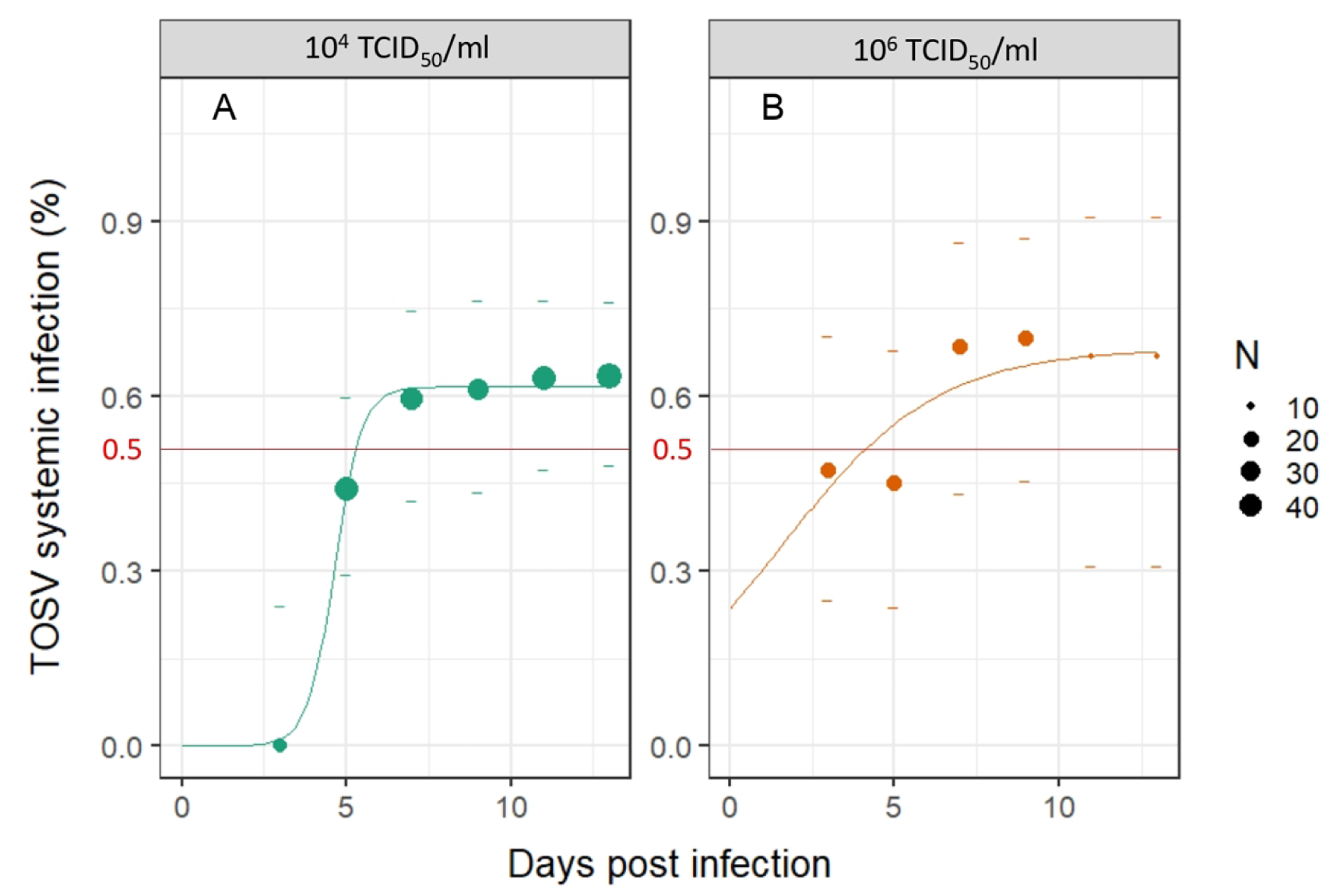
Systemic infection dynamics for *Ph. perniciosus* infected with 10^4^ TCID_50_/ml (A) and 10^6^ TCID_50_/ml (B). Cumulative rates of systemic (disseminated) infections over time post Toscana virus infection are represented as point, dot size indicates the number of samples. Dashes represent the 95% confidence intervals of rates, and the red line represents 50% systemic infection. The fitted values obtained with a non-linear regression model are represented by a line.

To determine the virus infectivity in sand fly samples, TOSV titers of bodies, heads, wings and legs were determined by an end-point dilution assay (TCID_50_). Four sand flies per day (every two days) were analyzed by titration for the highest infection (dose 3). Of the 84 samples analyzed (N = four sand flies * tree parts * seven days = 84), seven sand fly bodies were found positive: at 1, 3, 5, 7, 9, 11 and 14 dpi (**Table 2**). For each dpi, 1 body out of 4 sand fly samples was positive, representing a 25% infection rate. There were no positive samples for heads, legs and wings. Only the highest dose 3 was analyzed by titration for this experiment to ensure results were obtained, but it would also be necessary to further analyze samples from dose 2.

**Table 2.** Toscana virus titers (TCID_50_/ml) in *Phlebotomus perniciosus* bodies per day post infection (dpi) at dose 3 (10^6^ TCID_50_/ml).

| Dpi | 1 | 3 | 5 | 7 | 9 | 11 | 14 |
| --- | --- | --- | --- | --- | --- | --- | --- |
| TCID<sub>50</sub>/ml | 9.3 | 9.3 | 29 | 20 | 20 | 9.3 | 9280 |

Of the sugar cotton pads analyzed at two, four, six, eight and 10 dpi, in order to detect the virus released by the females during sugar feeding, TOSV was detected at both latest time points. According to the standard curve, the virus limit detection was 10^-2^ (log10) RNA copies/ml. An average of 10^3^ (log10) RNA copies/ml of TOSV was detected in samples collected at two dpi (30 female pool). At six (27 female pool) and eight (20 female pool) dpi, viral particles were detected in the sugar samples at an average of 10^3^ and 10^4^ (log10) RNA copies/ml, respectively.

### Toscana virus impact on vector life history traits

To investigate the impact of TOSV infection on *Ph. perniciosus* survival life-history trait, 140 and 98 sand flies were used for the control (uninfected females) and the test (infected females with dose 3), respectively. Blood-fed females were euthanized and dissected directly after oral infection. They ingested an average of 2.5x10^5^ RNA copies/ml. Using the Kaplan-Meier method, the survival distributions were not significantly different between the two groups (log-rank test, *P* = 0.92). The survival probabilities were not statistically different for uninfected and infected sand flies (S1 Appendix). The last infected female was purposely euthanized at 45 dpi, dissected and analyzed by TOSV RT-qPCR. An amount of 2.2x10^4^ RNA copies/ml was found in the body of this sample at 45 dpi. However, this data was considered censored for the survival analysis.

For the experimental investigation assessing the infection impact on vector fecundity life-history traits (eggs and larvae numbers, oviposition and hatching times), a cohort of 543 female specimens was used. The blood-feeding rates were relatively low for each group (50%). A total of 85 (control), 94 (dose 2) and 96 (dose 3) individuals were placed in individual laying pots. The different life-history trait data were collected during 35 days of monitoring of control and TOSV infected sand flies until the last hatching (**Table 3**). Among all the parameters studied, only hatching time of test groups dose 2 (GLM test, *P* = 0.0247) and dose 3 (GLM test, *P* = 5.61e-07) were significantly different from the control group. We have confirmed these significant results with the Kaplan-Meier method (**S2 Appendix**). Hatching time shows a positive correlation with the quantity of TOSV ingested (**Table 3**, **Fig 3**).

**Figure 3.**
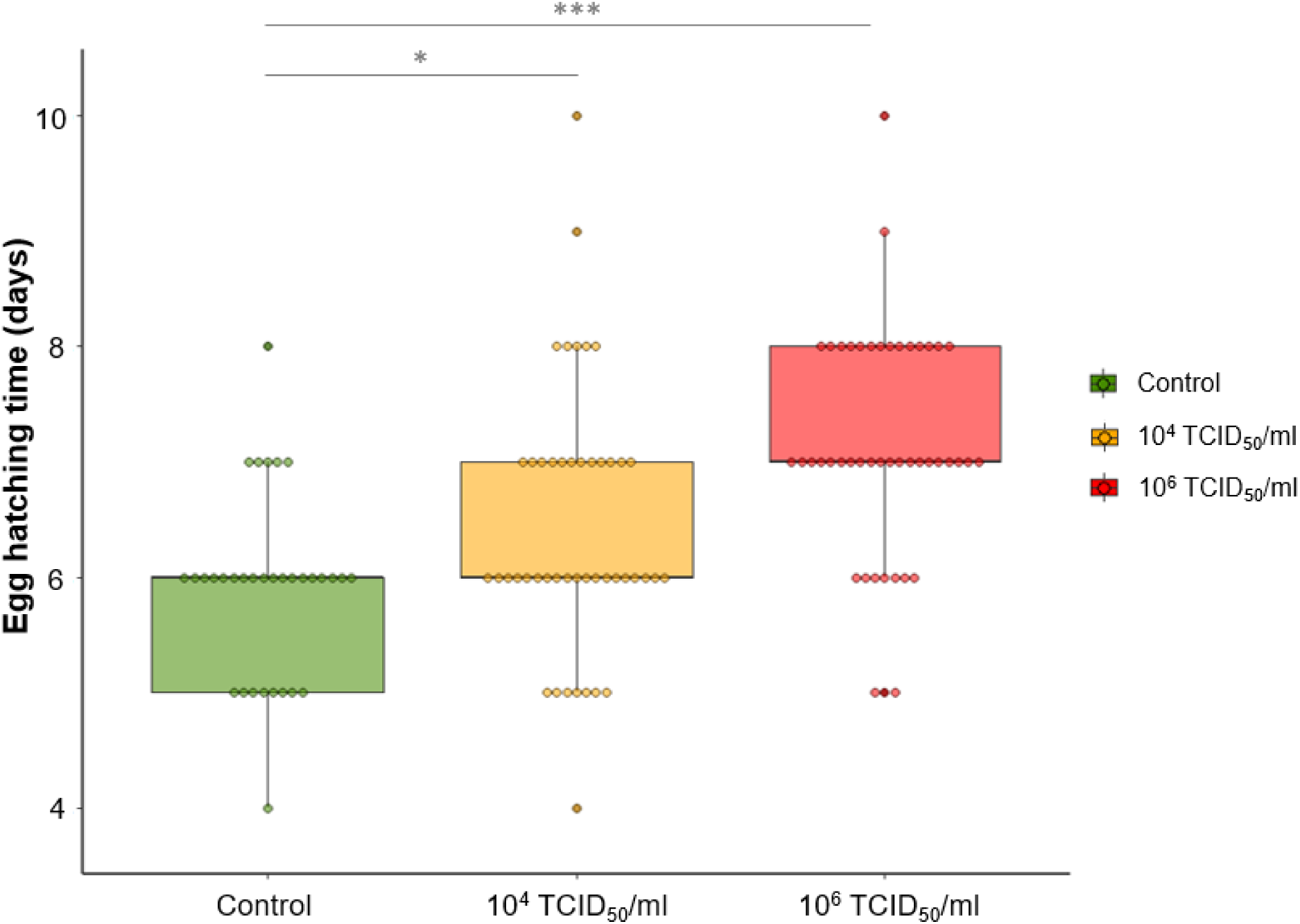
Boxplot representing the impact of Toscana virus infection on *Phlebotomus perniciosus* egg hatching time. Dots represent the average hatching time of eggs from one female. In green, the control group of uninfected females; in yellow, the test group of females infected with 10^4^ TCID_50_/ml; in red, the test group of females infected with 10^6^ TCID_50_/ml. The stars are the result of the GLM-based comparison (p-value significance: 0 ’***’ 0.001 ’**’ 0.01 ’*’ 0.05).

**Table 3.** Summary of the experiments with the different life history traits of *Phlebotomus perniciosus* studied. BF: Blood-Fed, BF*: Blood-Feeding, No: Number of. Means are indicated with their standard error.

|  | No. ♀<br>initial | No. ♀<br>BF | BF*<br>rate (%) | Survival<br>time (days) | No. eggs | Oviposition<br>time (days) | No. larvae | Hatching time<br>(days) |
| --- | --- | --- | --- | --- | --- | --- | --- | --- |
| Control | 170 | 85 | 50 | $7.6 \pm 0.4$ | $29.0 \pm 2.2$ | $6.0 \pm 0.4$ | $21.8 \pm 2.0$ | $5.9 \pm 0.1$ |
| Test $10^4$ | 182 | 94 | 52 | $9.2 \pm 0.3$ | $32.2 \pm 2.0$ | $7.9 \pm 0.4$ | $21.1 \pm 1.8$ | $6.4 \pm 0.2$ |
| Test $10^6$ | 191 | 96 | 50 | $8.6 \pm 0.4$ | $27.9 \pm 2.1$ | $7.2 \pm 0.3$ | $19.5 \pm 1.6$ | $7.1 \pm 0.1$ |

## Discussion

Toscana virus, an emerging sandfly-borne phlebovirus in the Mediterranean basin, is one of the most frequent cause of summer viral meningitis in geographic areas where human population are exposed and can lead to severe neurological cases (6). Despite a significant effect on human health, limited research has been conducted on its transmission dynamics. Although the main sand fly vectors of TOSV have been identified, the viral development cycle in its vectors remains largely unexplored (25). While numerous field studies have detected TOSV in *Ph. perniciosus* populations, only a few experimental studies have yet assessed the vectorial competence and capacity of this species (12,26,27). Considering this knowledge gap, our research focused on examining the dynamics of TOSV infection in *Ph. perniciosus* and its impact on various life history traits of this vector. By investigating these aspects, we aim to enhance our understanding of TOSV transmission and its effects on vector biology.

In our study, the first objective was to investigate experimentally infection dynamics in the vector, and to do this we used three different viral doses. In general, vector competence (intrinsic ability of a vector to transmit a virus) is a progressive process that begins when the vector takes an infectious blood meal from a viremic host. After initial infection and replication in the midgut, the virus spreads from the midgut to secondary tissues like haemocoel (systemic infection), then to the salivary glands, where it may be released in saliva during a subsequent blood meal (28). Herein, we found viral particles in head and thorax, where the salivary glands are presents (29), until at least 15 dpi, as well as in wings and legs (**Fig 1**). In addition, the viral titration in sand fly bodies infected with dose 3 (10^6^ TCID_50_/ml) showed that the TOSV was still infectious until 14 dpi (**Table 2**). In this study, we confirmed the TOSV replication and dissemination within the species *Ph. perniciosus*.

Additionally, our study demonstrated a direct correlation between the infection rates in sand flies and the infectious dose in the ingested blood meal (**Fig 1**). When sand flies were infected with a low TOSV dose (10^2^ TCID_50_/ml), virus particles did not disseminate in the vector, and only a few body samples were tested positive before sand fly defecation. Therefore, it can be assumed that infections at such low doses do not lead to systemic infection. In contrast, for sand flies infected with medium dose 2 (10^4^ TCID_50_/ml) and high dose 3 (10^6^ TCID_50_/ml) doses, TOSV disseminated into the secondary tissues and the infection persisted for at least 15 days (**Fig 2**). Based on our experimental findings, the minimum infectious dose for TOSV systemic infection in *Ph. perniciosus* is > 10^2^ TCID_50_/ml, although the exact threshold remains to be determined in a more detailed way using a specifically designed experimental protocol.

This result is in the line with other arbovirus model, as a study demonstrated that doses as low as 10^3.9^ plaque-forming units (PFU) per ml (approximately 10^4^ TCID_50_/ml) of an Asian genotype Chikungunya virus strain can lead to transmission by *Aedes albopictus* (30). Viremia levels in phlebovirus infections for human or non-human vertebrates, whether observed in the wild or the laboratory, are generally low and transient, often necessitating the ingestion of a substantial viral load by the vectors (12,31–33). In one of the few experimental infection studies involving *Ph. perniciosus* and TOSV lineage A, the rate of infected sand flies decreased over time despite ingestion of infectious doses greater than 10^6^ TCID_50_/ml (12). Conversely, our study showed that even when ingesting a medium dose of the virus (10^4^ TCID_50_/ml), the virus is detectable at least 21 days and the infected sand flies are therefore likely to transmit the virus. This discrepancy between the two studies may be due to the utilization of different TOSV lineages, which could influence vectorial competence and infection dynamics. After ingestion of a viremic blood meal, the viral load in a competent vector usually increases significantly to reach a plateau which is maintained for the duration of the vector’s life (34), which is well represented for TOSV as shown in **Fig 2**. We also detected TOSV RNA in the body of one sand fly at 45 dpi, suggesting that the vector can occasionally remain infected and might be infectious for a longer period. Since sand flies have a longevity of seven to 10 weeks (35), a specific experiment aiming at testing sand flies for a longer period (>45 dpi) would be necessary to determine whether the vector remains infectious. For several arboviruses, the viremia peak of symptomatic human infections is estimated to be around 10^6^ TCID_50_/ml (36,37). Nevertheless, asymptomatic human cases can achieve viremia levels as high as the lowest estimates for symptomatic cases (38). To date, no studies have reported the viral load of TOSV in asymptomatic patients. Our results showed that sand flies can become long-term infected with a lower dose (dose 2, 10^4^ TCID_50_/ml) than the viremia peak dose (dose 3, 10^6^ TCID_50_/ml). Therefore, sand flies could be infected throughout the viremic period, whether or not the person is symptomatic.

Subsequently, we explored the capacity of sand flies to transmit TOSV, specifically examining their ability to spit out the virus, contained in their saliva, during the feeding process (39). In the laboratory, female sand flies die before taking a second blood meal (40), precluding to measure transmission during a second artificial blood meal. Therefore, we employed an alternative to this issue, using the vector sugar feeding behavior to detect the virus (41). To predigest sugar, sand flies release saliva during feeding (42). We assumed that infected sand flies might be able to release viral particles in their saliva when they feed on sugar soaked cotton pads. As, expected, we confirmed that results observed with Massilia virus and *Ph. Perniciosus* (14) were similar with TOSV: viral RNA was detected in the sugar cotton pads at two, six, and eight dpi on which sand flies infected with high dose (10^6^ TCID_50_/ml) had fed. While the presence of TOSV particles at two dpi could be attributed to viral residues (contamination between individuals, defecation, blood residues in mouthparts, etc.), TOSV RNA at six and eight dpi may indicate its presence in the salivary glands. It has already been shown that *Culex spp*. mosquitoes release arboviruses during sugar feeding (43). Our results suggest that TOSV reaches the salivary glands of *Ph. perniciosus* between three and six dpi although this should be confirmed by fine dissection and testing of only the salivary glands. In addition, the release of TOSV into sugar meal together with recent demonstration of long-term persistence of infectious TOSV in sugar meal (up to 7 days at 26°C) (13) suggest that sand fly to sand fly transmission can occur orally during sap feeding in nature. To better understand the transmission dynamics and be able to predict TOSV emergence it would be crucial to investigate further the possible alternative transmission routes of TOSV between sand flies through their sugar meal.

Transmission becomes possible after the completion of the extrinsic incubation period (EIP), which is the interval between ingestion of virus and the earliest continued time at which virus is released in the saliva (34). The average EIP of sandfly-borne phleboviruses is at least one week (44). The EIP of the phlebovirus Rift Valley Fever virus (RVFV) is a minimum of 14 days after infection (45). As mentioned above, we hypothesized that TOSV reaches the vector salivary glands around six dpi. Based on these findings, we can postulate that TOSV EIP in *Ph. perniciosus* is approximately six days (**Fig 4**), which would be shorter than of other phleboviruses investigated (45,46). In addition, we showed the dose-effect on the systemic infection, occurring approximately at five and three dpi for sand flies infected by dose 2 (10^4^ TCID_50_/ml) and dose 3 (10^6^ TCID_50_/ml), respectively (**Fig 2**). Indeed, EIP can be influenced by several factors (*e.g.* environmental conditions, vector lifespan or immune system) including the ingested dose of virus (47–49). In order to accurately estimate the TOSV EIP in the vector, it would be interesting to carry out additional replicates of the sugar cotton pad test and further experiments to examine other parameters that may influence EIP. This parameter is essential since pathogens with a shorter EIP result in earlier transmission after infection (50).

**Figure 4.**
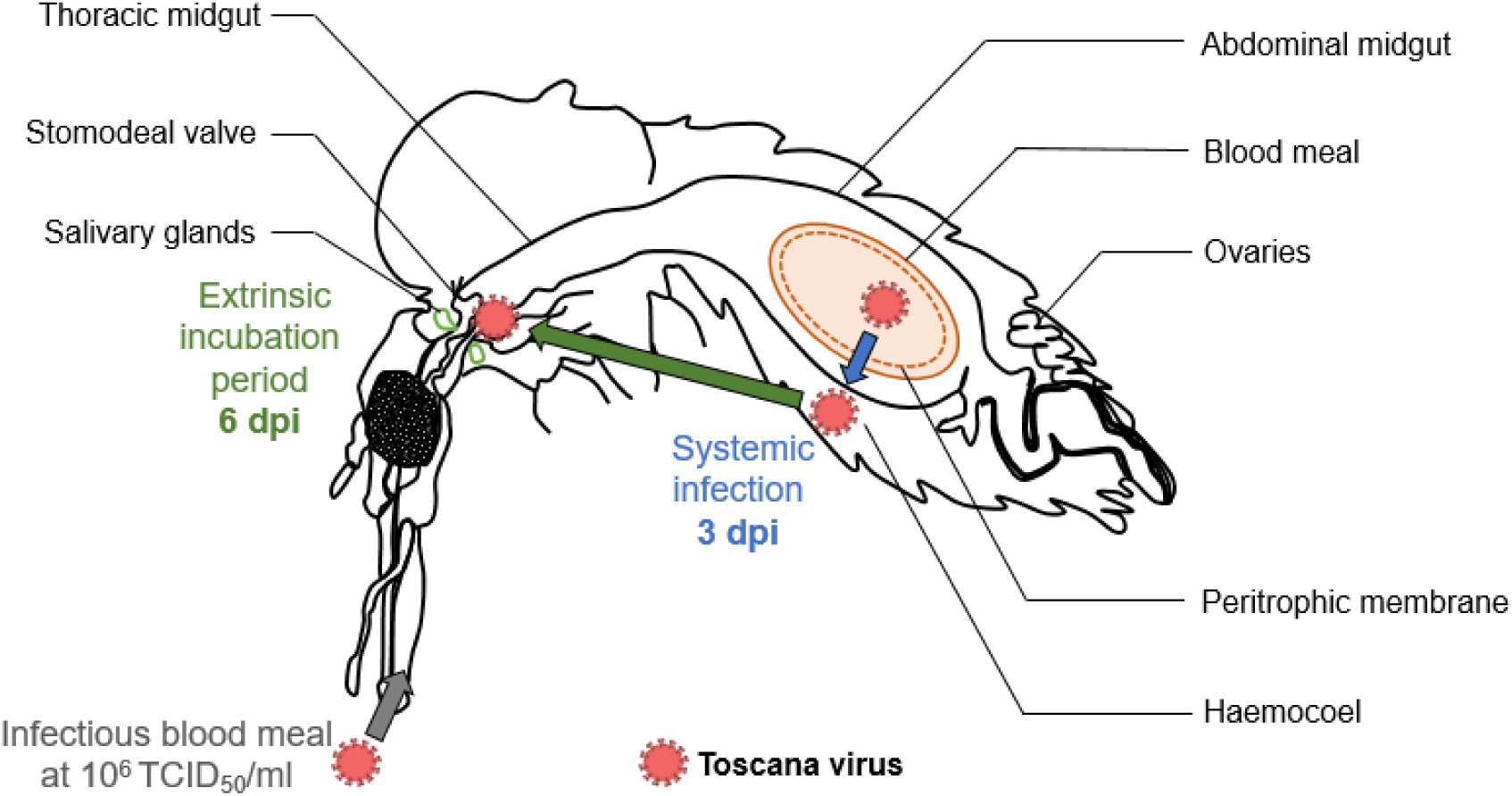
Schematic of the Toscana virus infection dynamics in the sand fly species *Phlebotomus perniciosus* including estimation of the extrinsic incubation period (green arrow) and the systemic infection (blue arrow) for the highest tested infectious dose 10^6^ TCID_50_/ml. dpi: days post infection

The second objective of our study was to explore the impact of TOSV infection on the sand fly biology, as effects on vector life-history traits are likely to have a strong impact on virus epidemiology. Initially, we sought to determine if high-dose TOSV infection could influence the vector’s survival. Our results indicated that the lifespan of infected sand flies was similar to that of uninfected ones (**S1 Appendix**). Therefore, it seems that high-dose TOSV infection have no impact on the vector survival. If the survival of infected sand fly remains unchanged, TOSV can be transmitted throughout the adult vector’s life during multiple blood meals (for females) and sugar meals (for males and females), as well as through multiple oviposition (vertical transmission), increasing the transmission risk. Then, we assessed the impact of two TOSV infection doses (10^4^ and 10^6^ TCID_50_/ml) on various life-history traits of *Ph. perniciosus* to determine the infection impact on vector biology. We showed the impact on the egg hatching time, which was longer for eggs from infected females (**Fig 3**). As the transovarial transmission of TOSV exists in *Ph. perniciosus*, we can hypothesize that infected sand flies will emerge later, potentially leading to an extension of the period of TOSV intra-species (sand flies) and inter-species (vertebrates and sand flies) transmission (51). A delay in vector emergence can lead to a reduction in intra- and inter-species competitiveness (52). We therefore propose an additional hypothesis whereby infected individuals that emerge later may have a better survival probability due to reduced competitiveness. Therefore, the next step should be to monitor the eggs of infected sand flies until adult emergence.

To conclude, we confirmed in this study that *Ph. perniciosus* is a competent vector for TOSV even at a lower dose in the blood meal (10^4^ TCID_50_/ml) than the virus dose during viremia peak in symptomatic mammals (10^6^ TCID_50_/ml). All our results suggest that the transmission of TOSV is short (six days) compared to other arboviruses (Phlebovirus (*e.g.* RVFV): 14 days (45), Flavivirus (*e.g.* ZIKV): 10 days approximately (47)) and that the epidemic risk in endemic areas must be considered. We have successfully developed and implemented effective protocols for conducting experimental infections with TOSV and *Ph. perniciosus*. We also demonstrated the TOSV infection impact on sand fly life history traits (egg hatching time) that could impact the virus epidemiology in natural populations. These experimental results provide a better understanding of virus maintenance in sand fly populations and the natural cycle of TOSV, while raising new hypotheses on transmission dynamics (**Fig 5**).

**Figure 5.**
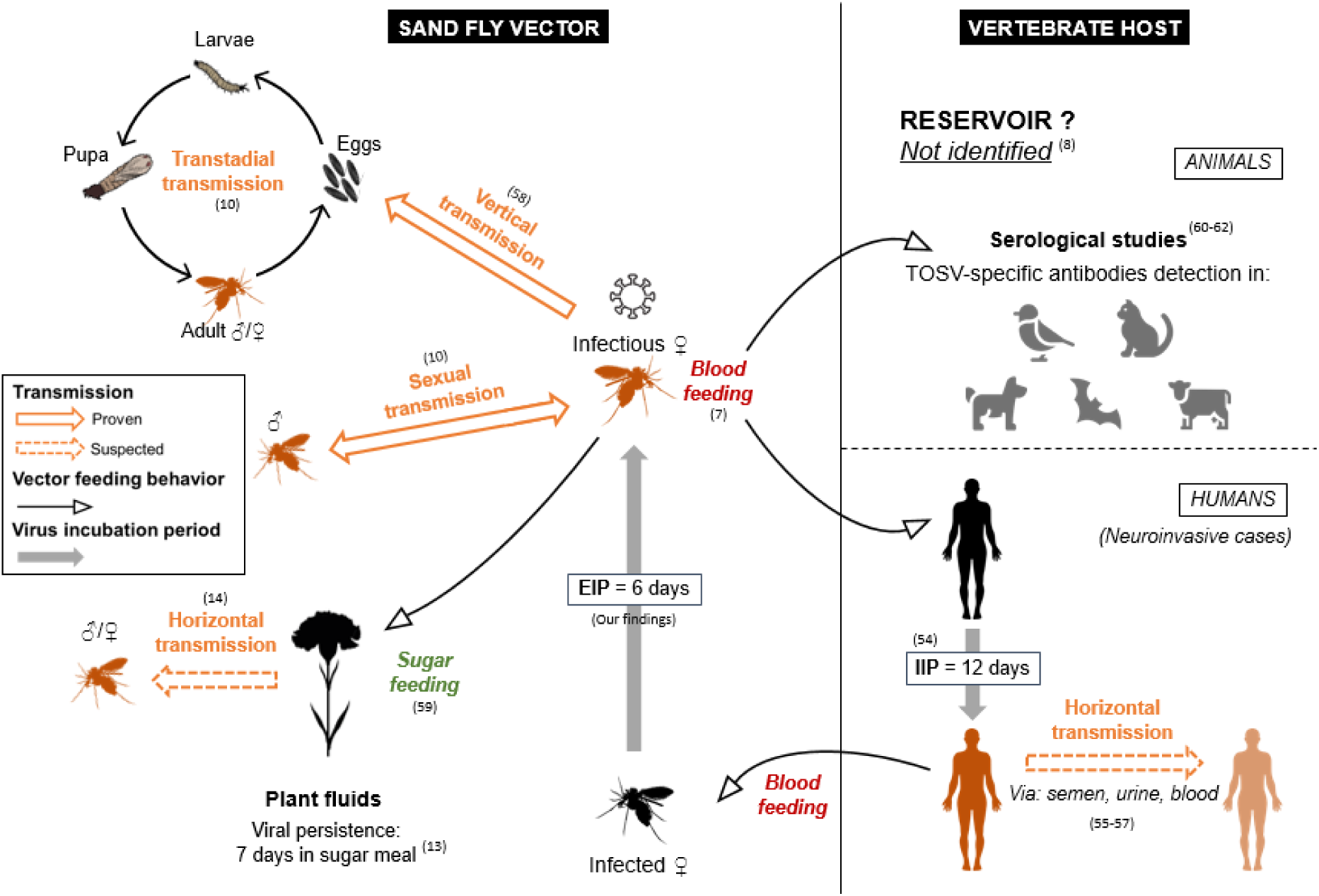
Toscana virus transmission dynamics within sand fly vectors and vertebrate hosts, including proven and suspected transmission routes, animals identified positive for TOSV-specific antibodies suspected but not identified as reservoirs, and incubation period estimates within the vector (EIP: Extrinsic Incubation Period) and the human neuroinvasive host (IIP: Intrinsic Incubation Period) (7,8,59–62,10,13,14,54–58).

## Materials and Methods

### 1. Sand fly rearing

In order to realize experimental studies, we established (IRD Vectopôle, Montpellier, France) a *Ph. perniciosus* (from Pr. Ricardo Molina, Laboratorio de Entomología Médica, Instituto de Salud Carlos III, Madrid, Spain) colony according to previously described protocols (63–65) with few modifications (40). Sand fly colonies were maintained in climate chamber under standard conditions (26 ± 1°C, 80% relative humidity (rh), 14h: 10h light: dark cycle). Adults were fed with cotton pads soaked in 50% organic sugar solution. Blood meal were realized once a week with glass feeders filled with rabbit blood maintained at 37°C and covered with a 3-day-old chick skin membrane. Feeding was performed for six hours with a regular manual blood mix to avoid clotting. After 24h, blood-fed females were transferred to plastic pots filled with a 1cm layer of plaster of Paris. These pots were stored in plastic boxes containing moistened sand. After hatching, larvae were fed with a macerated mixture of 40% rabbit excrement, 40% rabbit food and 20% mouse food. Newly emerged adults were transferred to rearing cages or in small cages for experimental infection.

### 2. Toscana virus experiments

#### Virus isolates

Lyophilized TOSV aliquots (strain MRS2010, lineage B) were provided by the laboratory UVE (Unité des Virus Emergents, Marseille, France). Vero E6 cells were grown in monolayers in Minimum Essential Medium (MEM, Gibco™) complemented with 7% heat-inactivated Fetal Bovin Serum (FBS, Eurobio Scientific), 1% L-glutamine (Gibco™) and 1% penicillin-streptomycin (Gibco™) at 37 °C with 5% CO_2_. Stocks of TOSV were obtained by dissolving the lyophilisates in pure water, and used for infecting Vero E6 cells at a multiplicity of infection (MOI) of 0.1 after five days of post infection. TOSV stocks at a concentration of 4.2 x 10^6^ 50% tissue culture infective dose (TCID_50_/ml) were aliquoted and stored at -80°C.

#### Experimental infection

Experimental infections of females *Ph. perniciosus* with TOSV were performed in enhanced biosafety level 3 (BSL-3) laboratory (IRD Vectopôle, Montpellier, France). Experimental infection were realized as previously described for *Leishmania* infection (66) modified for TOSV. Twenty-four hours prior to the experiment, 5 to 9-day-old females from the colony were starved by deprivation of sugar meal. Before infection, approximately 100 females were transferred into a feeding pot and stored in the BSL-3 climate chamber under standard conditions (26 ± 1°C, 80% rh) to acclimatize. Approximately 10% males were added to mate and stimulate females to take blood (63). Glass feeders were fill with heat-inactivated rabbit blood and covered with a chick skin membrane. For infected batches, TOSV was added at a final concentration of 10^2^ TCID_50_/ml (dose 1), 10^4^ TCID_50_/ml (dose 2) or 10^6^ TCID_50_/ml (dose 3), small, medium and large dose, respectively. For control batch (control), the same volume of MEM culture medium, without virus, was added. The complete system was set up in the climate chamber where the feeders were connected to a 37°C circulating water bath and fixed to the feeding pots (**Fig 6**). Feeding was performed for six hours, with reduced brightness, and regular manual blood mix to avoid clotting. Six hours of blood feeding was chosen as TOSV remains infectious and at the same amount in the blood meal under these conditions as previously described (13). As engorged females are fragile, it is recommended to handle them after the first 24 hours of digestion (63). Thus, they were sorted on ice the day after blood meal. Males and non-engorged females were killed and removed. All the experiments were replicated twice.

**Figure 6.**
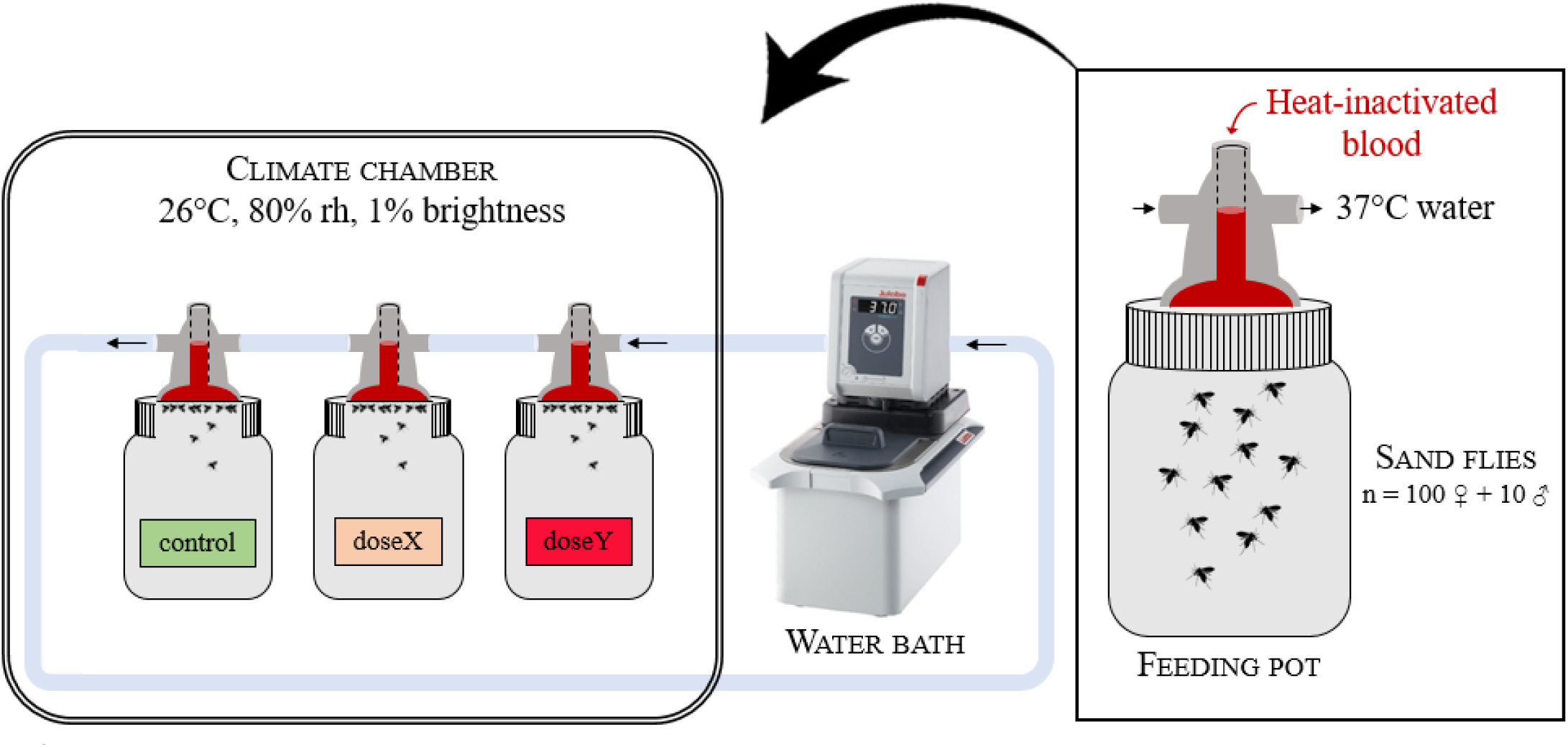
Blood feeding system in the BSL-3 laboratory. rh: relative humidity.

##### a. Infection dynamics

Engorged females were placed by 30 in cardboard boxes in climate chamber, under standard conditions, with a daily intake of cotton pads soaked in 50% organic sugar solution. Three groups were set up for this experiment, infected with dose 1, dose 2 and dose 3 of TOSV. Between 10 to 30 individuals were dissected in three parts: head, body (thorax and abdomen), wings and legs, every day for 15 days (**Fig 7**), and stored at -80°C for further TOSV detection and quantification analysis. In order to detect virus released by the females, the sugar cotton pads were collected daily, and stored at -80°C. One cotton per day at two, four, six, eight and 10 days post infection (dpi) with dose 3 were analyzed with the same RT-qPCR protocol as the sand fly samples. A standard curve was carried out for absolute quantification of TOSV RNA in sugar soaked cotton pads through RT-qPCR to obtain the virus detection limit (S3 Appendix).

**Figure 7.**
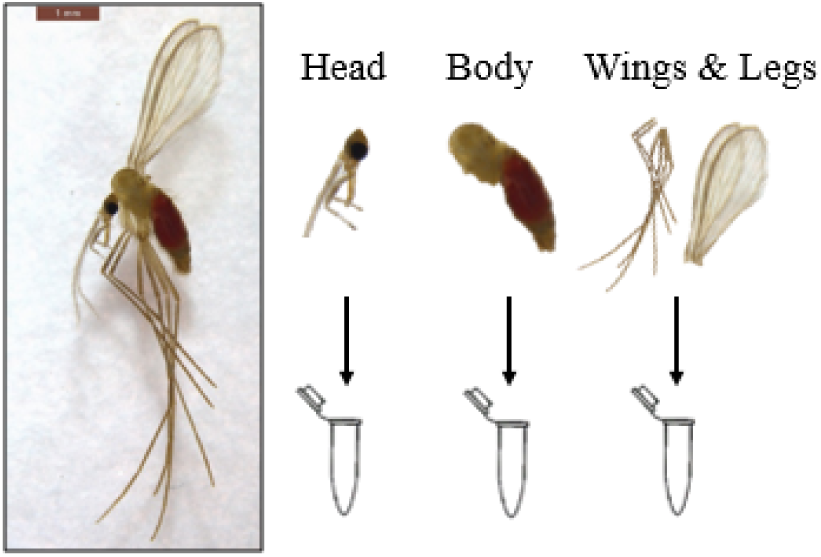
Dissection of infected females in three parts: head, body, wings and legs for Toscana virus detection (Photo by J. Prudhomme)

##### b. Impact of infection on life-history traits

To investigate the potential impact of TOSV infection on sand fly life-history traits, the traits most important in the virus transmission, which are vector survival and fecundity, were measured. For this experiment, engorged females were individually dispatched in small egg-laying pots filled with a 1cm layer of plaster of Paris, pooled by group in plastic boxes containing moistened sand and placed in climate chamber under standard conditions. A pool of sand flies was set aside in a cardboard box for TOSV infection verification by RT-qPCR. Groups were set up for this experiment, control with uninfected individuals, and two tests with individuals infected by different TOSV doses (10^4^ and 10^6^ TCID_50_/ml). Life history traits were measured daily and individually for each female: death date, oviposition date, eggs laid number, egg-hatching date and larvae number. Immature stages were only monitored up to first instar larva, due to potential cannibalism between larvae (67), and larvae were killed directly after hatching.

To measure infection impact on female survival, the experiment was performed with a different batch of females that did not have access to an egg-laying pot, and were therefore unable to lay eggs. Indeed, blood-fed females died quicker with the stress egg-laying induced by the rearing conditions (40). After blood feeding, engorged females were transferred to cardboard boxes with a maximum of 30 individuals per box in climate chamber and maintained under standard conditions. Five blood-fed females were killed, dissected and stored at -80°C for RT-qPCR analysis directly after oral infection to measure the initial amount of virus ingested. The survival time of two female groups, fed with (i) healthy blood (control) and (ii) infected blood with high dose of TOSV (dose3), was measured every day until the death of all individuals. In order to check the potential impact on survival, this experiment was only realized with the highest virus dose. To prevent the fungi development which could affect the survival of the sand flies, dead females were removed and the sugar source was changed every day.

#### Virus detection

Viral load was analyzed by RT-qPCR. Entire female or each part (head, body, wings and legs) were individually crushed in 300µl of MEM (enriched with 1% penicillin–streptomycin, 1% (200mM) l-glutamine, 1% kanamycin, and 3% amphotericin B (Gibco)). Sand fly tissues were homogenized by a Tissuelyser II (Qiagen) using 3mm tungsten carbide beads (Qiagen), and centrifuged at 10000 rpm for 5min. A volume of 200µl homogenate supernatant was used for nucleic acid extraction with the QIAcube (Qiagen) machine, using Virus Extraction Mini Kit (Qiagen). The RT-qPCR assay were performed with a SuperScript III Platinum One-Step RT-qPCR Kit with ROX (Invitrogen - Thermo Fisher Scientific) on a QuantStudio 12K Flex thermocycler (ThermoFisher). A volume of 5µl of RNA was added to 20µl of mix containing 12.5µl of 2X Reaction Mix, 0.5µl of Superscript III RT/Platinum Taq Mix, and primers and probes STOS at 10µM (68). Negative (pure water) and positive controls (at 4.81 x 10^4^ RNA in vitro transcribed copies/ml as described by Beckert & Masquida (69)) were included in each RT-qPCR run; samples with Ct value < 40 were considered positive.

TOSV titers were determined by end-point dilution assay. Tenfold dilutions were used to infect confluent Vero E6 cells in a 96-well plate in MEM (5% FBS, 1% penicillin-streptomycin, 1% kanamycin, 3% amphotericin B (Gibco)) at 37°C and 5% CO_2_. Wells were classified as positive (cytopathic effect) *versus* negative (no cytopathic effect) at five dpi and TCID_50_/ml was calculated according to the Reed-Muench method (70).

### 3. Statistical analysis

Data analysis were carried out with R through Rstudio software version 2022.07.1 - © 2009-2022 (71). Survival was examined with the Kaplan Meier product limit estimator, and a log-rank test was used to compare survival distributions between uninfected and TOSV-infected sand flies, using the packages survival (72) and survminer (73). Egg hatching times were also examined with the Kaplan Meier method, and a log-rank test was used to compare hatching time distributions between uninfected and TOSV-infected sand flies with both doses (10^4^ and 10^6^ TCID_50_/ml). The significance of the TOSV doses (10^4^ and 10^6^ TCID_50_/ml) and the effect of time post-infection on the infection dynamics was tested by general linear models (GLM). Probabilities of systemic infection according to the TOSV dose or/and time post-infection were estimated with a logistic model. Median systemic infection doses were calculated based on logistic regression parameter estimates. **Fig 2** was made using the package ggplot2 (74).

## Acknowledgments

Authors thank Dr. Justine Boutry (IRD, Montpellier, France) and Dr. Marc Choisy (OUCRU, Hanoi, Vietnam) for their valuable assistance in the statistical analysis; Justine Fournier, Aloïs Berard (Montpellier University, France) and Dr. Idris Mhaidi (IRD, Montpellier, France) for their help to maintain the sand fly colonies; and Dr. Maarten Schrama (Institute of Environmental Sciences, Leiden University, Leiden, Netherlands) for reviewing the manuscript and providing valuable feedback.

## Author contributions

L.L., A.-L.B. and J.P. conceptualized the study. The study design method was discussed with L.L., J.P., A.-L.B., N.A., R.C., and A.F. The experiments were carried out by L.L. with support from N.A. and J.P. Also, L.L. analyzed the data, following models created by A.F., and L.L. wrote the manuscript with the support of J.P., A.-L.B., N.A., R.C., and A.F. Finally, L.L., J.P., A.-L. B., N.A., R.C., and A.F. read, amended and approved the final version of the manuscript.

## Supporting information

**S1 Appendix.**
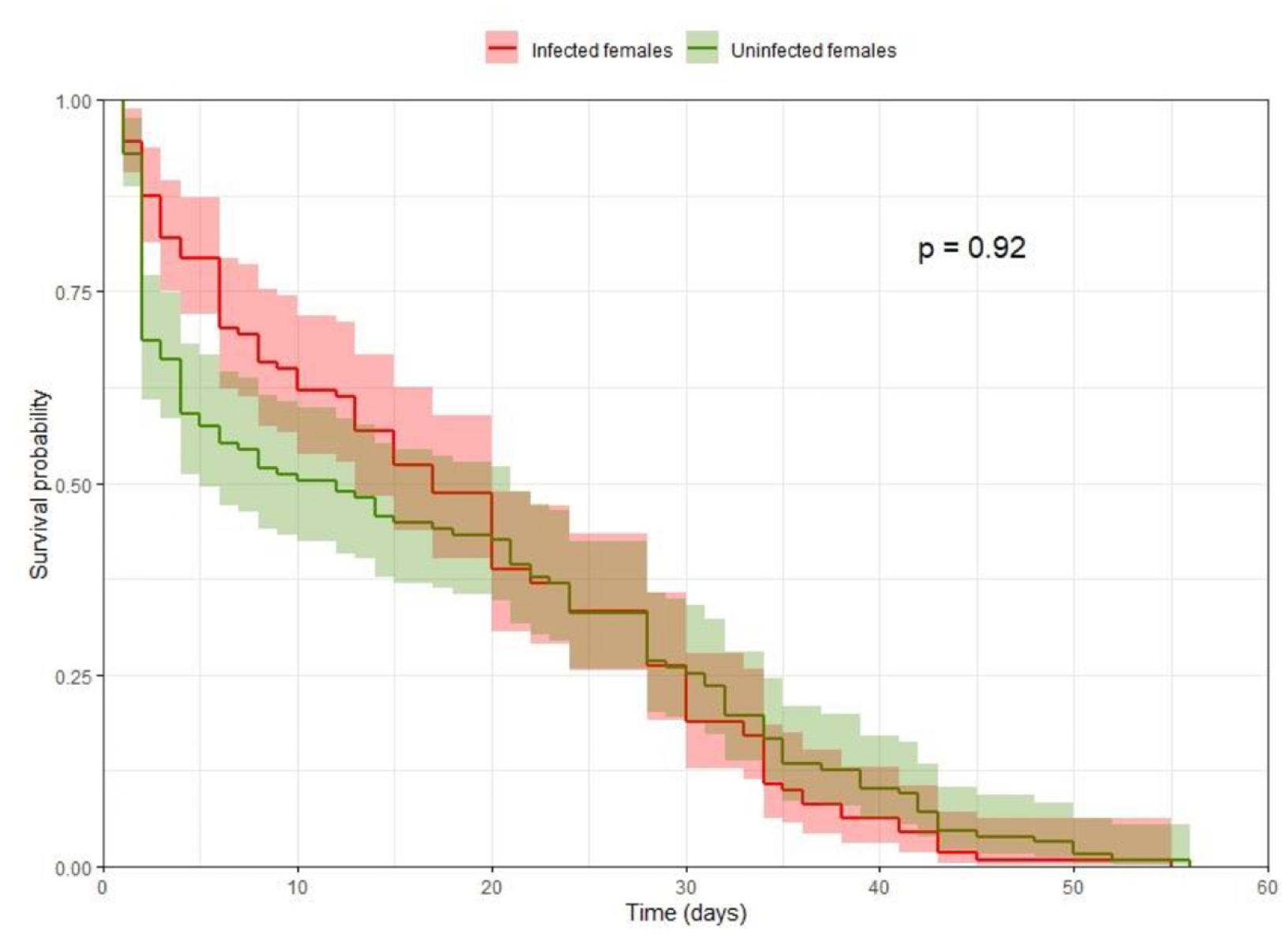
Kaplan-Meier survival estimates for infected in red (n=98) and uninfected in green (n=140) sand flies with Toscana virus with 95% confidence intervals

**S2 Appendix.**
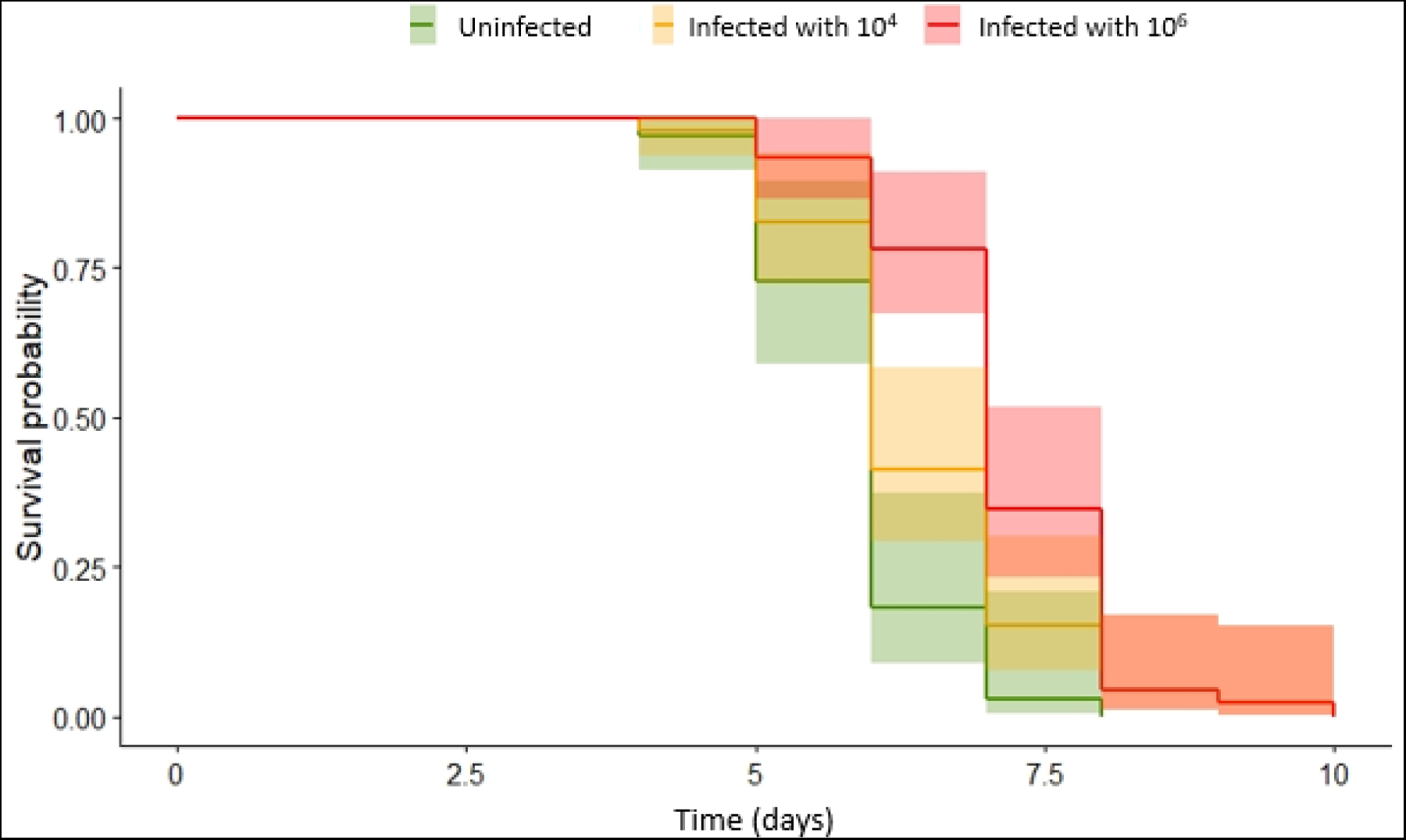
Kaplan-Meier hatching time estimates for Toscana virus infected in orange (10^4^ TCID_50_/ml, n=62) and red (10^6^ TCID_50_/ml, n=62); and uninfected in green (n=46) sand flies with 95% confidence intervals

**S3 Appendix.**
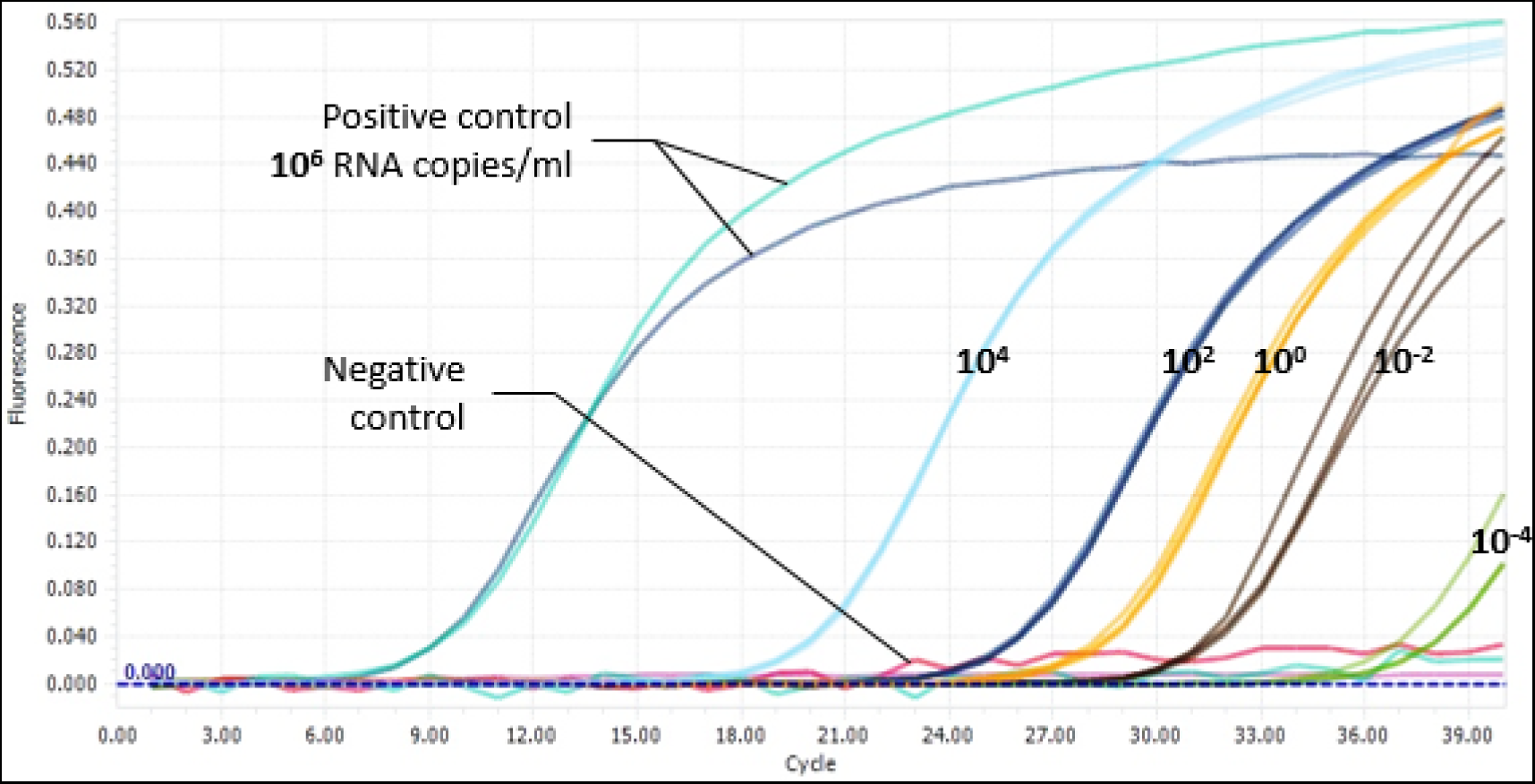
Standard curve for absolute quantification of the Toscana virus RNA in sugar solution by real-time reverse transcriptase-polymerase chain reaction. All the samples were amplified and considered positive, except samples at 10^-4^ RNA copies/ml considered as negative because of late amplification (>36 cycles). So the virus detection limit is 10^-2^ RNA copies/ml in the sugar solution.

## References

1. Kuhn JH, Abe J, Adkins S, Alkhovsky S V., Avšič-Županc T, Ayllón MA, et al. Annual (2023) taxonomic update of RNA-directed RNA polymerase-encoding negative-sense RNA viruses (realm Riboviria: kingdom Orthornavirae: phylum Negarnaviricota). J Gen Virol [Internet]. 2023 Aug 25 [cited 2024 Jan 10];104(8):001864. Available from: https://www.microbiologyresearch.org/content/journal/jgv/10.1099/jgv.0.001864

2. Verani P, Ciufolini MG, Caciolli S, Renzi A, Nicoletti L, Sabatinelli G, et al. Ecology of viruses isolated from sand flies in Italy and characterized of a new *Phlebovirus* (Arabia virus). Am J Trop Med Hyg [Internet]. 1988 [cited 2022 Sep 30];38(2):433–9. Available from: https://pubmed.ncbi.nlm.nih.gov/3128131/

3. ECDC 2022. Toscana virus infection [Internet]. [cited 2022 Nov 14]. Available from: https://www.ecdc.europa.eu/en/toscana-virus-infection

4. Charrel RN, Bichaud L, de Lamballerie X. Emergence of Toscana virus in the mediterranean area. World J Virol. 2012;1(5):135.

5. Braito A, Corbisiero R, Corradini S, Marchi B, Sancasciani N, Fiorentini C, et al. Evidence of Toscana virus infections without central nervous system involvement: A serological study. Eur J Epidemiol. 1997;13(7):761–4.

6. Charrel RN, Gallian P, Navarro-Marí JM, Nicoletti L, Papa A, Sánchez-Seco MP, et al. Emergence of Toscana virus in Europe. Emerg Infect Dis. 2005;11(11):1657–63.

7. Depaquit J, Grandadam M, Fouque F, Andry PE, Peyrefitte C. Arthropod-borne viruses transmitted by Phlebotomine sandflies in Europe: A review. Eurosurveillance. 2010;15(10):40–7.

8. Ayhan N, Prudhomme J, Laroche L, Bañuls AL, Charrel RN. Broader geographical distribution of toscana virus in the mediterranean region suggests the existence of larger varieties of sand fly vectors. Microorganisms. 2020;8(1):e114.

9. Ayhan N, Charrel RN. Of phlebotomines (sandflies) and viruses: a comprehensive perspective on a complex situation. Curr Opin INSECT Sci. 2017;22:117–24.

10. Tesh RB, Lubroth J, Guzman H. Simulation of arbovirus overwintering: Survival of Toscana virus (Bunyaviridae:Phlebovirus) in its natural sand fly vector *Phlebotomus perniciosus*. Am J Trop Med Hyg. 1992;

11. Ciurolini MG, Maroli M, Guandalini E, Marchi A, Verani P. Experimental studies on the maintenance of Toscana and Arbia viruses (Bunyaviridae: *Phlebovirus*). Am J Trop Med Hyg [Internet]. 1989 Jun 1 [cited 2022 May 17];40(6):669–75. Available from: https://europepmc.org/article/med/2545100

12. Ciufolini MG, Maroli M, Verani P. Growth of two phleboviruses after experimental infection of their suspected sand fly vector, Phlebotomus perniciosus (Diptera: Psychodidae). Am J Trop Med Hyg [Internet]. 1985 Jan 1 [cited 2022 Sep 27];34(1):174–9. Available from: https://europepmc.org/article/med/3918472

13. Laroche L, Ayhan N, Charrel R, Bañuls A-L, Prudhomme J. Persistence of Toscana virus in sugar and blood meals of phlebotomine sand flies: epidemiological and experimental consequences. Sci Reports 2023 131 [Internet]. 2023 Apr 5 [cited 2023 Apr 11];13(1):1–7. Available from: https://www.nature.com/articles/s41598-023-32431-9

14. Jancarova M, Bichaud L, Hlavacova J, Priet S, Ayhan N, Spitzova T, et al. Experimental infection of sand flies by massilia virus and viral transmission by co-feeding on sugar meal. Viruses. 2019 Apr 1;11(4):1–15.

15. Wilder-Smith A, Gubler DJ, Weaver SC, Monath TP, Heymann DL, Scott TW. Epidemic arboviral diseases: priorities for research and public health. Lancet Infect Dis. 2017 Mar 1;17(3):e101–6.

16. Heitmann A, Jansen S, Lühken R, Leggewie M, Badusche M, Pluskota B, et al. Experimental transmission of zika virus by mosquitoes from central Europe. Eurosurveillance [Internet]. 2017 Jan 12 [cited 2022 Sep 23];22(2):30437. Available from: https://www.eurosurveillance.org/content/10.2807/1560-7917.ES.2017.22.2.30437/

17. Soni M, Khan SA, Bhattacharjee CK, Dutta P. Experimental study of dengue virus infection in *Aedes aegypti* and *Aedes albopictus*: A comparative analysis on susceptibility, virus transmission and reproductive success. J Invertebr Pathol. 2020 Sep 1;175:107445.

18. Coffey LL, Failloux AB, Weaver SC. Chikungunya virus–vector interactions. Viruses. 2014;6(11):4628–63.

19. Tabachnick WJ. Ecological effects on arbovirus-mosquito cycles of transmission. Curr Opin Virol. 2016 Dec 1;21:124–31.

20. Lequime S, Dehecq JS, Matheus S, Laval F De, Almeras L, Briolant S, et al. Modeling intra-mosquito dynamics of Zika virus and its dose-dependence confirms the low epidemic potential of *Aedes albopictus*. PLOS Pathog [Internet]. 2020 Dec 31 [cited 2022 Sep 23];16(12):e1009068. Available from: https://journals.plos.org/plospathogens/article?id=10.1371/journal.ppat.1009068

21. Kramer LD, Ciota AT. Dissecting vectorial capacity for mosquito-borne viruses. Curr Opin Virol. 2015 Dec 1;15:112–8.

22. Styer LM, Meola MA, Kramer LD. West nile virus infection decreases fecundity of Culex tarsalis females. J Med Entomol. 2007;44(6).

23. da Silveira ID, Petersen MT, Sylvestre G, Garcia GA, David MR, Pavan MG, et al. Zika Virus Infection Produces a Reduction on Aedes aegypti Lifespan but No Effects on Mosquito Fecundity and Oviposition Success. Front Microbiol. 2018;9.

24. Costanzo KS, Muturi EJ, Montgomery A V., Alto BW. Effect of oral infection of La Crosse virus on survival and fecundity of native Ochlerotatus triseriatus and invasive Stegomyia albopicta. Med Vet Entomol. 2014;28(1).

25. P Verani, MG Ciufolini, L Nicoletti, M Balducci, G Sabatinelli, M Coluzzi, P Paci LA. Ecological and epidemiological studies of Toscana virus, an arbovirus isolated from *Phlebotomus*. Ann Ist Super Sanita. 1982;18(3):397–9.

26. Daoudi M, Calzolari M, Boussaa S, Bonilauri P, Torri D, Romeo G, et al. Identification of Toscana virus in natural population of sand flies (Diptera: Psychodidae) from Moroccan leishmaniasis foci. J Infect Public Health. 2022 Apr 1;15(4):406–11.

27. Ayhan N, Alten B, Ivovic V, Martinkovic F, Kasap OE, Ozbel Y, et al. Cocirculation of Two Lineages of Toscana Virus in Croatia. Front Public Heal. 2017;5(December):1– 4.

28. Fontaine A, Jiolle D, Moltini-Conclois I, Lequime S, Lambrechts L. Excretion of dengue virus RNA by *Aedes aegypti* allows non-destructive monitoring of viral dissemination in individual mosquitoes. Sci Rep [Internet]. 2016;6(February):1–10. Available from: 10.1038/srep24885

29. Nacif-Pimenta R, Pinto LC, Volfova V, Volf P, Pimenta PFP, Secundino NFC. Conserved and distinct morphological aspects of the salivary glands of sand fly vectors of leishmaniasis: An anatomical and ultrastructural study. Parasites and Vectors [Internet]. 2020 Sep 3 [cited 2023 Oct 5];13(1):1–12. Available from: https://parasitesandvectors.biomedcentral.com/articles/10.1186/s13071-020-04311-y

30. Ledermann JP, Borland EM, Powers AM. Minimum infectious dose for chikungunya virus in *Aedes aegypti* and *Ae. albopictus* mosquitoes. Rev Panam Salud Pública. 2017 Aug 21;41:e65.

31. Bartelloni PJ, Tesh RB. Clinical and serologic responses of volunteers infected with phlebotomus fever virus (Sicilian type). Am J Trop Med Hyg [Internet]. 1976 May 1 [cited 2022 Sep 27];25(3):456–62. Available from: https://europepmc.org/article/med/180844

32. Tesh RB, Duboise SM. Viremia and immune response with sequential phlebovirus infections. Am J Trop Med Hyg [Internet]. 1987 May 1 [cited 2022 Sep 27];36(3):662–8. Available from: https://europepmc.org/article/med/3034088

33. Grazia Cusi M, Gori Savellini G, Terrosi C, Di Genova G, Valassina M, Valentini M, et al. Development of a mouse model for the study of Toscana virus pathogenesis. Virology. 2005 Mar 1;333(1):66–73.

34. Mellor PS. Replication of arboviruses in insect vectors. Vol. 123, Journal of Comparative Pathology. W.B. Saunders Ltd; 2000. p. 231–47.

35. Oerther S, Jöst H, Heitmann A, Lühken R, Krüger A, Steinhausen I, et al. Phlebotomine sand flies in Southwest Germany: An update with records in new locations. Parasites and Vectors [Internet]. 2020 Apr 21 [cited 2023 Oct 6];13(1):1–8. Available from: https://parasitesandvectors.biomedcentral.com/articles/10.1186/s13071-020-04058-6

36. Tsetsarkin KA, Vanlandingham DL, McGee CE, Higgs S. A single mutation in Chikungunya virus affects vector specificity and epidemic potential. PLoS Pathog. 2007;3(12):1895–906.

37. Mcintosh BM, Paterson HE, Donaldson JM, De Sousa J. Chikungunya Virus: Viral Susceptibility and Transmission Studies with some Vertebrates and Mosquitoes. S Afr J Med Sci. 1963;28(112):45–52.

38. Chastel C. Eventual role of asymptomatic cases of dengue for the introduction and spread of dengue viruses in non-endemic regions. Front Physiol. 2012;3 MAR:70.

39. Schneider CA, Calvo E, Peterson KE. Arboviruses: How Saliva Impacts the Journey from Vector to Host. Int J Mol Sci 2021, Vol 22, Page 9173 [Internet]. 2021 Aug 25 [cited 2023 Oct 6];22(17):9173. Available from: https://www.mdpi.com/1422-0067/22/17/9173/htm

40. Prudhomme J, Laroche L, Nguyen Hoang Quan A, Fournier J, Mhaidi I, Berard A, et al. Establishment and maintenance of two sand fly colonies (*Phlebotomus perniciosus* and *Phlebotomus papatasi*) for experimental infections. Submitt to Jounal Med Entomol. 2022;

41. Hall-Mendelin S, Ritchie SA, Johansen CA, Zborowski P, Cortis G, Dandridge S, et al. Exploiting mosquito sugar feeding to detect mosquito-borne pathogens. Proc Natl Acad Sci U S A [Internet]. 2010 Jun 22 [cited 2023 Nov 15];107(25):11255–9. Available from: https://www.pnas.org/doi/abs/10.1073/pnas.1002040107

42. Charlab R, Valenzuela JG, Rowton ED, Ribeiro JMC. Toward an understanding of the biochemical and pharmacological complexity of the saliva of a hematophagous sand fly *Lutzomyia longipalpis*. Proc Natl Acad Sci [Internet]. 1999 Dec 21 [cited 2022 Sep 27];96(26):15155–60. Available from: https://www.pnas.org/doi/abs/10.1073/pnas.96.26.15155

43. Hurk AF van den, Johnson PH, Hall-Mendelin S, Northill JA, Simmons RJ, Jansen CC, et al. Expectoration of Flaviviruses During Sugar Feeding by Mosquitoes (Diptera: Culicidae). J Med Entomol [Internet]. 2007 Sep 1 [cited 2022 Mar 29];44(5):845–50. Available from: https://academic.oup.com/jme/article/44/5/845/973141

44. Brett-Major DM, Claborn DM. Sand Fly Fever: What Have We Learned in One Hundred Years? Mil Med [Internet]. 2009 Apr 1 [cited 2022 Aug 18];174(4):426–31. Available from: https://academic.oup.com/milmed/article/174/4/426/4333798

45. Moutailler S, Bouloy M, Failloux A-B. Efficient oral infection of *Culex pipiens quinquefasciatus* by Rift Valley fever virus using a cotton stick support. Am J Trop Med Hyg. 2007;76(5):827–9.

46. Yuko E, Sang R, Owino EA, Ingonga J, Matoke-Muhia D, Hassaballa IB, et al. Sandfly Blood-Feeding Habits and Competence in Transmitting Ntepes Virus, a Recently Discovered Member of the Genus *Phlebovirus*. Biomed Res Int. 2022;2022.

47. Winokur OC, Main BJ, Nicholson J, Barker CM. Impact of temperature on the extrinsic incubation period of Zika virus in *Aedes aegypti*. PLoS Negl Trop Dis [Internet]. 2020 Mar 1 [cited 2022 Sep 28];14(3):e0008047. Available from: https://journals.plos.org/plosntds/article?id=10.1371/journal.pntd.0008047

48. Anderson SL, Richards SL, Tabachnick WJ, Smartt CT. Effects of West Nile virus dose and extrinsic incubation temperature on temporal progression of vector competence in *Culex pipiens quinquefasciatus*. J Am Mosq Control Assoc [Internet]. 2010 Mar [cited 2022 Sep 28];26(1):103. Available from: /pmc/articles/PMC2858365/

49. Mbaika S, Lutomiah J, Chepkorir E, Mulwa F, Khayeka-Wandabwa C, Tigoi C, et al. Vector competence of *Aedes aegypti* in transmitting Chikungunya virus: Effects and implications of extrinsic incubation temperature on dissemination and infection rates. Virol J [Internet]. 2016 Jun 29 [cited 2022 Sep 28];13(1):1–9. Available from: https://virologyj.biomedcentral.com/articles/10.1186/s12985-016-0566-7

50. Bellan SE, Cornell SJ. The Importance of Age Dependent Mortality and the Extrinsic Incubation Period in Models of Mosquito-Borne Disease Transmission and Control. PLoS One [Internet]. 2010 [cited 2022 Sep 28];5(4):e10165. Available from: https://journals.plos.org/plosone/article?id=10.1371/journal.pone.0010165

51. Tesh RB, Modi GB. Maintenance of Toscana virus in *Phlebotomus perniciosus* by vertical transmission. Am J Trop Med Hyg. 1987;36(1):189–93.

52. Leisnham PT, Juliano SA. Impacts of climate, land use, and biological invasion on the ecology of immature Aedes mosquitoes: implications for La Crosse emergence. Ecohealth [Internet]. 2012 Jun [cited 2023 Dec 5];9(2):217–28. Available from: https://pubmed.ncbi.nlm.nih.gov/22692799/

53. Navarro-Mari JM, Palop-Borras B, Perez-Ruiz M, Sanbonmatsu-Gamez S. Serosurvey study of Toscana virus in domestic animals, Granada, Spain. Vector Borne Zoonotic Dis. 2011 May;11(5):583–7.

54. Laroche L, Jourdain F, Ayhan N, Bañuls AL, Charrel R, Prudhomme J. Incubation Period for Neuroinvasive Toscana Virus Infections. Emerg Infect Dis [Internet]. 2021 Dec 1 [cited 2022 Feb 28];27(12):3147. Available from: /pmc/articles/PMC8632186/

55. Matusali G, D’Abramo A, Terrosi C, Carletti F, Colavita F, Vairo F, et al. Infectious Toscana Virus in Seminal Fluid of Young Man Returning from Elba Island, Italy. Emerg Infect Dis [Internet]. 2022 Apr 1 [cited 2022 Sep 26];28(4):865. Available from: /pmc/articles/PMC8962903/

56. Ergunay K, Kaplan B, Okar S, Akkutay-Yoldar Z, Kurne A, Arsava EM, et al. Urinary detection of toscana virus nucleic acids in neuroinvasive infections. J Clin Virol. 2015;

57. Brisbarre N, Attoui H, Gallian P, Bonito P Di, Giorgi C. Seroprevalence of Toscana virus in blood donors, France, 2007. Emerg Infect Dis [Internet]. 2011 [cited 2022 Mar 28];17(5):941–3. Available from: https://stacks.cdc.gov/view/cdc/24349/cdc_24349_DS1.pdf

58. Maroli M, Ciufolini MG, Verani P. Vertical transmission of Toscana virus in the sandfly, Phlebotomus perniciosus, via the second gonotrophic cycle. Med Vet Entomol [Internet]. 1993 Jul 1 [cited 2023 Nov 17];7(3):283–6. Available from: https://onlinelibrary.wiley.com/doi/full/10.1111/j.1365-2915.1993.tb00689.x

59. Molyneux DH, Moore J, Maroli M. Sugars in sandflies. Parassitologia [Internet]. 1991 Dec 1 [cited 2023 Nov 17];33 Suppl:431–6. Available from: https://europepmc.org/article/med/1841240

60. Ayhan N, López-Roig M, Monastiri A, Charrel RN, Serra-Cobo J. Seroprevalence of Toscana Virus and Sandfly Fever Sicilian Virus in European Bat Colonies Measured Using a Neutralization Test. Viruses 2021, Vol 13, Page 88 [Internet]. 2021 Jan 11 [cited 2023 Nov 17];13(1):88. Available from: https://www.mdpi.com/1999-4915/13/1/88/htm

61. Ayhan N, Rodríguez-Teijeiro JD, López-Roig M, Vinyoles D, Ferreres JA, Monastiri A, et al. High rates of antibodies against Toscana and Sicilian phleboviruses in common quail Coturnix coturnix birds. Front Microbiol. 2023 Jan 4;13:5056.

62. Alwassouf S, Maia C, Ayhan N, Coimbra M, Cristovao JM, Richet H, et al. Neutralization-based seroprevalence of toscana virus and sandfly fever sicilian virus in dogs and cats from Portugal. J Gen Virol [Internet]. 2016 Nov 1 [cited 2022 Feb 28];97(11):2816–23. Available from: https://www.microbiologyresearch.org/content/journal/jgv/10.1099/jgv.0.000592

63. Volf P, Volfova V. Establishment and maintenance of sand fly colonies. J Vector Ecol. 2011;36(SUPPL.1):1–9.

64. Lawyer P, Killick-Kendrick M, Rowland T, Rowton E, Volf P. Laboratory colonization and mass rearing of phlebotomine sand flies (Diptera, Psychodidae). Parasite. 2017;24.

65. Molina R, Jiménez MY, Alvar J. Methods in sand fly research. 2017. 108 p.

66. Vaselek S, Prudhomme J, Myskova J, Lestinova T, Spitzova T, Bañuls A-L, et al. Comparative Study of Promastigote-and Amastigote-Initiated Infection of *Leishmania infantum* (Kinetoplastida: Trypanosomatidae) in *Phlebotomus perniciosus* (Diptera: Psychodidae) Conducted in Different Biosafety Level Laboratories. J Med Entomol [Internet]. [cited 2020 Mar 3];57(2):601–7. Available from: http://creativecommons.

67. Srinivasan R, Panicker KN. Cannibalism among immatures of *Phlebotomus papatasi* (Diptera: Psychodidae). J Bombay Nat Hist Soc. 1992;89:386–7.

68. Perez-Ruiz M, Collao X, Navarro-Mari J-M, Tenorio A. Reversetranscription, real-time PCR assay for detection of Toscana virus. J Clin Virol. 2007 Aug;39(4):276–81.

69. Beckert B, Masquida B. Synthesis of RNA by In Vitro Transcription BT - RNA: Methods and Protocols. Methods Mol Biol [Internet]. 2011 [cited 2022 Jul 29];703:29–41. Available from: http://www.ncbi.nlm.nih.gov/pubmed/21125481

70. Reed LJ, Muench H. A Simple Method of Estimating Fifty Percent Endpoints. Am J Trop Med Hyg. 1938;27(20):493–7.

71. R devlopment Core Team. R: A Language and Environment for Statistical Computing. Vienna, Austria. 2019;

72. Therneau T. A Package for Survival Analysis in R [Internet]. 2022. p. R package version 3.4-0. Available from: https://cran.r-project.org/package=survival

73. Kassambara A, Kosinski M, Biecek P. Package survminer: Drawing Survival Curves using “ggplot2.” [Internet]. 2019. Available from: https://cran.r-project.org/package=survminer

74. Wickham H. ggpolt2: Elegant Graphics for Data Analysis. Use R! Ser [Internet]. 2016 [cited 2022 Sep 13];211. Available from: http://had.co.nz/ggplot2/book

